# Towards a Gamete Matching Platform: Using Immunogenetics and Artificial Intelligence to Predict Recurrent Miscarriage

**DOI:** 10.1101/534594

**Authors:** Aldo Mora-Sánchez, Daniel-Isui Aguilar-Salvador, Izabela Nowak

## Abstract

The degree of Allele sharing of the Human Leukocyte Antigen (HLA) genes has been linked with recurrent miscarriage (RM). However, no clear genetic markers of RM have yet been identified, possibly because of the complexity of interactions between paternal and maternal genes. We propose a methodology to analyse HLA haplotypes from couples either with histories of successful pregnancies or RM. This article describes, for the first time, a method of RM genetic-risk calculation. Novel HLA representation techniques allowed us to create an algorithm (IMMATCH) to retrospectively predict RM with an AUC = 0.71 (p = 0.0035) thanks to high-resolution typing and the use of linear algebra on peptide binding affinity data. The algorithm features an adjustable threshold to increase either sensitivity or specificity. Combining immunogenetics with artificial intelligence could create personalized tools to better understand the genetic causes of unexplained infertility and a gamete matching platform that could increase pregnancy success rates.

---

Women using assisted reproductive technologies have a significantly higher risk of experiencing miscarriage than those attempting natural pregnancies, even after adjusting for age ^1^. The cause is not identified for about 50% of patients who experience recurrent miscarriage (RM) ^2^. As identifying the cause has proven difficult, most preventing efforts are primarily focused on assessing the quality of gametes prior to fertilization ^3^ and quality of embryos before implantation ^4^. Currently, there are no methods to accurately assess an individual couple’s genetic risk of RM; such a risk prediction method would likely improve the process of gamete donation.

Immune interactions are proposed as a possible explanation for RM, as developing embryos can be considered as semi-allografts to the maternal immune system. The fact that women with autoimmune disorders have an increased incidence of RM lends credence to this theory ^5^. Among others, Human Leukocyte Antigen (HLA) proteins and Killer-cell immunoglobulin-like receptors (KIR) are proteins thought to mediate immune interactions in pregnancy.

HLA molecules are membrane proteins that present antigenic peptides to T cells, which makes them determinant molecules of the adaptive human immune system. HLA proteins are encoded by different genes and are one of the most polymorphic regions of the human genome ^6^. The most common explanation for this high variability is that it resulted from the competition between slowly evolving vertebrates and rapidly evolving pathogens ^7^. The set of HLA proteins differ from person to person, and each HLA variant has the ability to bind to different repertoires of peptides; this variation consequently determines the immune system’s ability to react to different pathogens. As each HLA variant is potentially able to act against different sets of pathogens, natural selection is hypothesized to favour heterozygous individuals ^8^. Both animal and human studies ^9^ suggest that mating behaviour favours HLA-diverse or HLA-dissimilar individuals. Indeed, couples more dissimilar in their HLA alleles have been reported to be more fertile ^10^. Although the specific signalling mechanisms responsible for HLA-mediated mating behaviour remains unclear, it is known that paternal and maternal haplotypes are not randomly matched genetically ^8,9^. In spite of that, this information is not part of the selection criteria when an assisted reproductive therapy patient receives or selects a gamete from an unknown external party, and therefore the matching is random at the genetic level.

KIRs, the second family of molecules thought to be involved in immune interactions during pregnancy, are membrane proteins expressed mainly by natural killer cells and by subsets of T cells ^11^. HLA Class I (HLA-A, -B and C) proteins interact with KIR, which act as activating or inhibitory receptors that regulate the effector properties of the cells that express them. Every individual can carry multiple and diverse copies of both activating and inhibitory receptors in their genome, creating a highly diverse genetic region.

The interactions between immunity and pregnancy outcomes have been demonstrated empirically at the population level, with several studies showing consistently that HLA allele sharing correlates with pregnancy outcomes ^12^. However, identifying specific genetic markers, especially for individual couples, remains elusive. Having observed the limitations present in previous studies, we propose two strategies to identify genetic markers.

Cognizant that HLA proteins are highly polymorphic, the first proposed strategy is to increase the resolution of their representation. The high degree of polymorphism implies that the potential number of combinations of paternal and maternal protein variants is extremely high, which poses serious difficulties for any study design. We can illustrate this high variability by examining only the three genes coding for HLA class I proteins: to date there are 3,172 reported HLA-A proteins, 3,923 HLA-B proteins, and 2,920 HLA-C proteins (http://hla.alleles.org/nomenclature/stats.html). For any given couple, the number of possible combinations is therefore 3,172^4^ + 3,923^4^ + 2,920^4^, more than 50,000 times the current earth’s population. No study design thus can make use of full resolution HLA genotyping for analysing the effect of HLA protein combinations on pregnancy. Regarding mathematical representations, HLA proteins are categorical variables with high cardinality (number of possible proteins); most researchers so far have dealt with this high cardinality by representing protein variants with ultra-low resolution. The study of genetic interactions often relies on dividing thousands of proteins into two functional groups. For instance, the joint effect of HLA-C and KIR haplotypes is usually investigated by categorizing HLA-C genes as either in the C1 or the C2 group, and KIR genes as homozygous or not for the A haplotype ^13^. In addition, allele sharing in research is commonly reduced to a similar/dissimilar relationship ^14^. As these assessments are binary, most of the information on diversity, key for HLA functioning, is lost. As increasing the resolution of HLA proteins would render certain analyses impossible, we propose feasible mathematical representation techniques to handle the high resolution information of HLA variability.

The second proposed strategy to identify individual RM genetic risk markers involves addressing the complexity of genetic interactions, as the joint effect of multiple genes is rarely considered in the literature. When studying the impact of maternal-paternal interactions on pregnancy outcomes, the vast majority of studies focus either on the effect of allele sharing at a single locus ^14,^, or on the interaction of two loci ^13^. Recent advances in machine learning techniques applied to medicine have showed their potential to analyze the joint effect of multiple variables and to consider their interactions. For a recent review of how machine learning can contribute to comprehensive, inexpensive, and accurate diagnostics, see ^15^.

The study described in this article is the first of its kind to simultaneously increase protein resolution while at the same time addressing multiple complex genetic interactions. We first constructed explanatory variables (features) to train our machine learning model to predict cases of RM. This process of constructing relevant features is denominated “feature engineering”. To our knowledge, our approach is the first of its kind in two aspects concerning feature engineering. First, we expressed the degree of allele sharing between paternal-maternal genes in a continuous, instead of binary, manner. Second, in the context of genetic associations, we used linear algebra methods to numerically represent HLA proteins by using peptide binding affinity data. The engineered features were used as the input of machine learning algorithms. Our method has several additional advantages over existing methods, specifically the fact that it is able to go beyond finding relevant predictors at the population level and is, to our knowledge, the first one able to predict probabilities on a case-by-case basis. The latter could be particularly advantageous for clinical applications, in particular personalized genetic counselling.

## RESULTS

### Discriminative power and accuracy

The classifier achieved an AUC = 0.71 (*p* = 0.0035). The AUC is a measure of the predictive power of the classifier, being AUC = 1 a perfect classifier and 0.5 a random classifier. The accuracy of the model was 0.67 (*p* = 0.0045).

### Specificity and sensitivity of the classifier

Accurately identifying the group a potential gamete pair belongs to could reduce the risk of RM in a gamete donation program. To this end, we calculated the specificity and sensitivity of the classifier (Figure 1). We can observe that it is possible to achieve a sensitivity of 86% if we accept a false positive rate of 57%. The clinical implications of accepting a false positive rate of 57% in the specific case of gamete donation are developed in the discussion.

**Figure 1.**
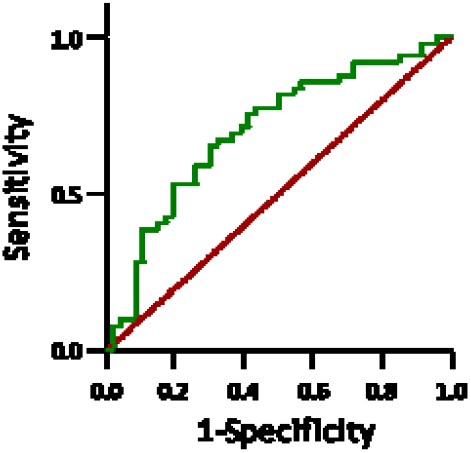
ROC curve, in which sensitivity and 1-specificity are plotted as a function of the classification threshold. A unique sensitivity-specificity pair exists for each threshold value. The green line corresponds to all possible sensitivity-specificity pairs obtained by varying the classifier’s threshold, whereas the red line corresponds to the sensitivity-specificity pairs of a random classifier.

Figure 2 shows how the classifier behaves when features are gradually added to the model, illustrating how the joint analysis of features outperforms the analysis of single features. Figure 3 compares the AUC of Immunomatch algorithm with that of a classifier trained with HLA-C1/C2 and KIR-A homozygosity information, as this interaction has been reported to be relevant at the population level for spontaneous abortion ^16^. In addition, the comparison is extended to the AUC of the feature that, alone, obtained the highest AUC value.

**Figure 2.**
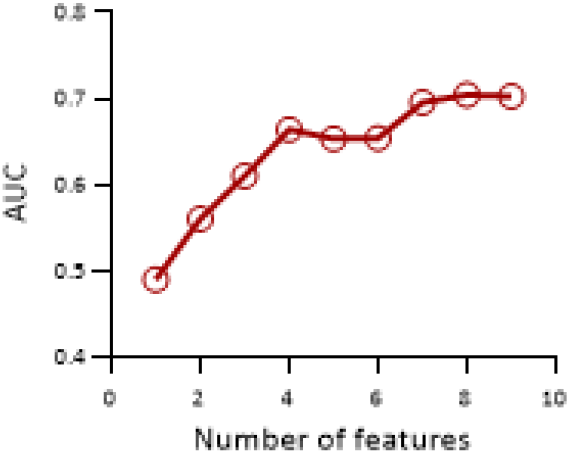
Classification performance measured as the AUC as a function of the number of features used to build the classifier, to evaluate the joint predictive power of features.

**Figure 3.**
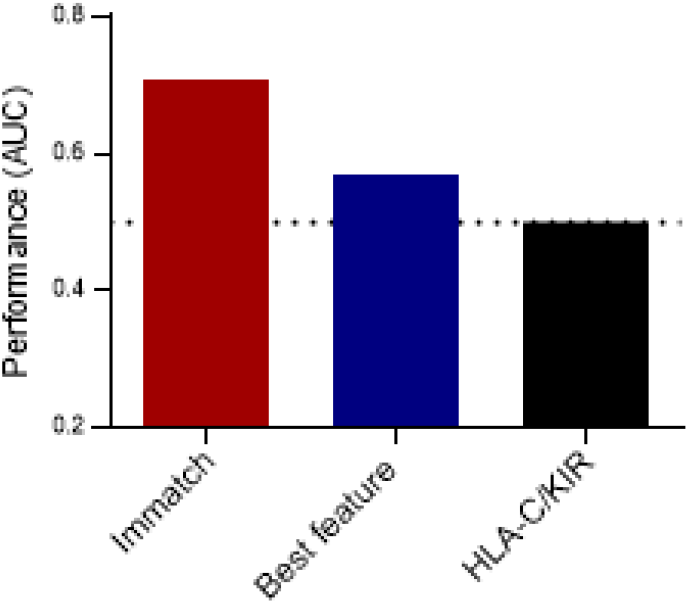
Performance of the algorithm (IMMATCH) compared to a classifier trained only with the best feature, and a classifier trained with a marker reported in the literature (HLA-C, KIR interactions). Dotted line indicates the performance of a random

## DISCUSSION

Incorporating the joint effect of predictors is the only way to analyse the underlying complexity of genetic interactions. In our study, single features performed no better than random chance in predicting the risk group membership of an individual case. The classifier using the reported interactions between HLA-C2 and KIR AA performed no better than random chance either. This illustrates how a reduction in haplotype resolution by dividing HLA-C and KIR genes into two groups (similar and dissimilar), a strategy that according to some evidence could lead to a relevant predictor of RM at the population level, might not be sufficient for predicting individual outcomes.

Accurately assessing the risk for obstetric complications associated with a given pair of gametes could improve gamete donor selection. Our classifier has an adjustable classification threshold value that can be raised or lowered to improve sensitivity or specificity. A low threshold would increase sensitivity at the expense of specificity. This scenario is particularly interesting for gamete donor selection, where sensitivity is undoubtedly more important than specificity. This difference is because the patient is expected to choose only one or two donors from a catalog whose size can range from dozens to thousands. With a sensitivity of 86% and a false positive rate of 57%, even if slightly more than half of the donors proposed as potentially leading to RM for a specific recipient are false negatives, there is still a 86% probability that the selected donor(s) will lead to a healthy pregnancy. The above situation contrasts with the case of a disease diagnostic test for an individual. In such case, false positives are as important as false negatives, since a false positive is likely to unnecessarily change the lifestyle of the patient.

Ultimately, and especially for gamete catalogues with a small number of donors, there is no need to divide the donors into low and high risk groups. Instead of setting a threshold and making two categories, donors could be ranked according to their likelihood of leading to a healthy pregnancy and this ranking would change from recipient to recipient, which would become a valuable additional criterion for recipients.

When more data becomes available, other types of feature selection techniques or classifiers better suited to large data sets might allow for the discovery of associations involving other genes. Furthermore, the physicochemical continuous representation of the proteins (see Methods section and the Supplementary Information) can be used itself as a set of features. Such representation would allow mapping the large number of different protein variants to a reduced number of continuous variables, rendering feasible the mathematical analysis of paternal-maternal genetic combinations.

It has been suggested that the immunogenetics field is not sufficiently advanced to guide clinical interventions for the prevention of negative pregnancy outcomes ^17^. One of the limitations for this is that very large studies are required to find genetic markers. It has been estimated that in order to detect the risk of reduced birth weight with 90% power, data from almost 4,000 pregnancies would be required. Although this estimated volume of data may be true for that particular problem with traditional statistics. Taking this problem as a classification task significantly reduces the required number of data needed. Taking the present study as an example, we can use the binomial distribution to compute the probability of correctly identifying, by chance, more than 67% of the 95 cases (49 belonging to one class and 46 to the other class). Such probability is as low as 0.13%. This corresponds to a p value of p=0.0013, consistent with our numeric estimation of the p-value via a permutation test. Therefore, even relatively small sample sizes can lead to significant *p* values in a classification task, as the probability of obtaining positive results by chance decreases exponentially with the number of items to be classified. Conversely, when using population statistics, if multiple hypotheses are tested, several associations with seemingly significant p-values may appear by chance. Traditional statistical tests require an additional p-value correction that incorporates the number of hypothesis; however, the output of a classifier is a single score that combines all the predictors into one hypothesis and does not require further corrections.

In summary, the feature extraction methods developed here allow us to take better advantage of the high-resolution data available through current sequencing technologies. The computed *p* value suggests that, when taken together, the proposed measures are indeed markers of RM, and the obtained classifier could predict RM on a case-by case basis. These results could be further extended to embryo selection, in combination with current screening techniques. It is unlikely that a handful of immune genes can completely explain a phenomenon as complex as RM, and therefore an AUC of 1 might not be realistic. However, larger data collection is likely to further increase the predictive power of the classifier. Continued research on this subject may help to unveil the genetic factors of unexplained infertility and further contribute to helping couples with difficulty conceiving.

## METHODS

### Study design

A subset of 200 DNA samples from a previous study on spontaneous abortion ^18^ was used. Patients and their partners were recruited from the Department of Surgical, Endoscopic and Oncologic Gynaecology and from the Department of Gynaecology and Gynaecologic Oncology from the Polish Mothers’ Memorial Hospital–Research Institute in Poland. The RM group was made up of patients who had experienced at least 3 spontaneous miscarriages, but had no prior history of chromosomal aberrations, uterine anomalies, or hormonal disturbances; no Toxoplasma, Chlamydia, Listeria, or Brucella infection; and who tested negative for the presence of autoantibodies. Only samples from couples where the female was under 37 years old were considered for this study. Given the inclusion criteria, the RM group was composed of cases of unexplained RM.

The control group was recruited from Department of Obstetrics and Gynaecology, Medical University of Warsaw and from the Strzelce Opolskie District Hospital. This group consisted of healthy couples with at least two children born healthy and no history of miscarriage or endocrinological or immunological disorders. Experimental protocols were approved by the local ethics committees (with the agreement of the Medical University of Wrocław and the Polish Mothers’ Memorial Hospital–Research Institute in Łódź) and informed consent was obtained from both members of the couples included in the study.

The 200 DNA samples corresponded to 100 couples. 50 of these couples belonged to the RM group and the remaining 50 couples belonged to the control group. 5 couples were not included because DNA concentration from 5 participants was not high enough to perform high resolution genotyping. The final number of couples analysed was therefore 95, with 49 RM examples and 46 healthy controls.

### DNA Preparation and Genotyping

Genomic DNA was isolated from venous blood using the Invisorb Spin Blood Midi Kit (Invitek, Berlin, Germany), following the manufacturer’s instructions. DNA was stored at −80°C until further use. DKMS Life Science Lab performed the HLA-A,B,C genotyping using their in-house Next Generation Sequencing methodology, accredited by the European Federation for Immunogenetics and the American Society for Histocompatibility and Immunogenetics. When necessary, amino acid sequence was inferred from the typing information using the IPD and IMGT/HLA Database ^19^.

### Feature extraction

We chose to express allele sharing in a continuous manner to go beyond the binary assessment of similarity/dissimilarity commonly used in the literature. Representing allele sharing with a continuous, rather than binary, value expresses the complete spectrum of similarity.

We propose using functional, structural and evolutionary properties of the HLA proteins to build functions that map two proteins into a continuous distance metric. In particular, we propose three such functions per locus, f_1_, f_2_ y f_3_, each of which leads to a variable, or feature. As there are two maternal (say, L_1_ and L_2_) and two paternal (L_3_ and L_4_) alleles at a locus L, each feature F_i_(L) is constructed as

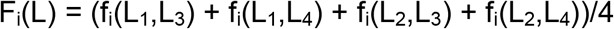

Each function f_i_ is meant to be a way to assess in a non-binary manner the extent to which two proteins are similar. Therefore, each feature F_i_ estimates, on average, how similar paternal and maternal proteins are at a given locus when using the function f_i_ for similarity assessment. The total number of features per couple is thus three times the number of loci considered.

The functions f_1_ and f_2_ assess the similarity, or distance, between two proteins in terms of peptide sequence similarity. Such assessment is performed via a substitution matrix, that estimates how similar two peptides are. The first function (f_1_) uses the BLOSUM50 ^20^ matrix (see Supplementary Information); the second function (f2) uses the PMBEC matrix ^21^.

The third function (f_3_) was derived using a novel technique that maps a protein to a vector of *n* numbers, by means of experimental measures regarding the binding affinity between that protein and reference peptides. These vectors, that are meant to describe physicochemical properties of the protein, are embedded in an n-dimensional abstract mathematical space. We will refer to this numeric representation as the *physicochemical continuous representation* of a protein. The similarity between two proteins is then computed as the inner product of the vectors representing the two proteins, as discussed in depth in the Supplementary Information. As the set of numbers conforming the vector represents physicochemical properties of the protein, we can expect this similarity assessment to provide information that is not redundant with the f_1_ and f_2_ assessments. The protein mapping was inspired by matrix factorization techniques ^22^ used for recommender systems, which are systems that use historical data to predict user preferences of certain items ^23^. We drew analogies between HLA proteins and users, peptides and items, and binding affinities and user preferences.

We decided to focus our analysis on allele sharing, as allele sharing consistently showed correlation with RM in the literature ^12^. Furthermore, we used only class I HLA genes for two reasons. The first reason is that f_3_ requires binding affinity data, which are not available for all of the class II HLA genes. The second reason is that, as diversity is key for the efficacy of all the HLA loci, any mechanism favouring it is likely to be observed across loci. Therefore, the genetic dissimilarity in a few HLA genes might be a proxy measure of the genetic dissimilarity of the remaining HLA genes. If, due to parsimony, a few genes must be chosen for the analysis, it is more convenient to choose the three class I genes, as they are more polymorphic than the class II HLA genes.

Specifically, for each couple 9 features were extracted, 3 features for each of the three HLA class I loci. Not enough binding affinity was available for 31 of the 1140 (six copies of each of the 190 participants) HLA proteins, and in these cases, the proteins were replaced by the closest ones as discussed in detail in the Supplementary Information.

### Classification

Classification refers to the process of predicting the class (in this case, either RM or healthy pregnancy) to which an example (the features of the previous section, derived from the genotype of a couple) belongs, using an algorithm trained on previous data (the features derived from the genotype of other couples) with a known class. An algorithm that predicts a certain example’s class membership is known as a classifier. We used a support vector machine (SVM) with a linear kernel as classifier for our study. This choice was done due to the sample size. For the same reason, to avoid overfitting, we tested only the default value of the margin constant C (C=1), the only hyperparameter of an SVM with a linear kernel. A hyperparameter is a parameter defined before the learning process, and therefore it is independent of the data. Further discussion on SVM hyperparameters can be found in ^24^. The process that starts with genotyping a couple and ends with predicting the probability to belong to the RM group is illustrated in Figure 4.

**Figure 4.**
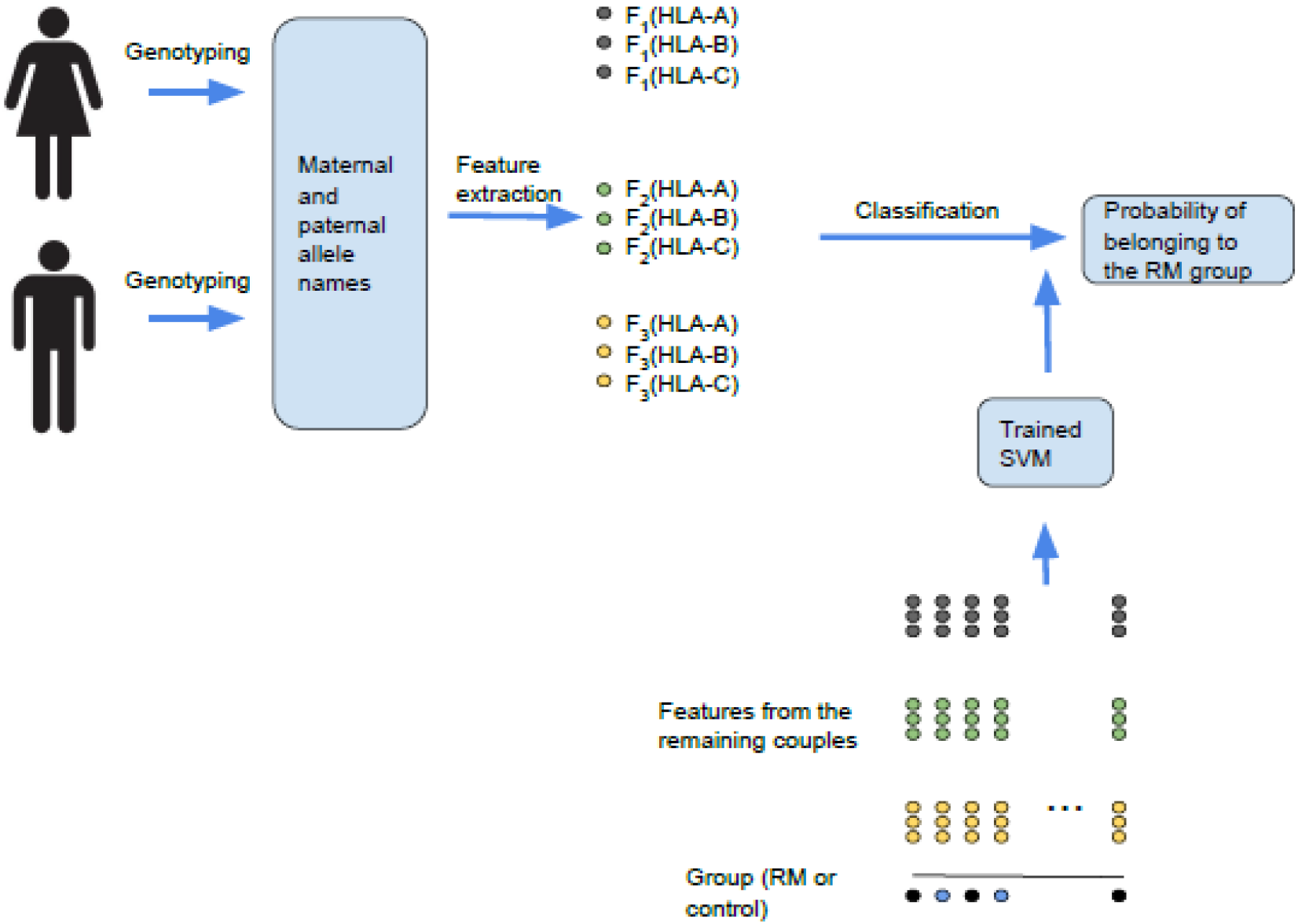
The classification process, from genotyping data to the estimation of the probability of belonging to the RM group. Leave-one-out cross validation iterates the above process for each couple. At each iteration, the selected couple is removed from the training set to evaluate the performance of the classifier with new data.

Leave-one-out cross-validation (LOOCV) was used to test the classifier’s ability to correctly predict the class to which new data belong ^25^. Broadly, LOOCV consists of removing one example, training the classifier with the remaining examples, predicting the class of the removed example, evaluating how accurate the prediction was, and finally computing the mean value of the accuracy by iterating over all possible examples. For N examples, LOOCV allows an independent test set of an effective size of N, and a training set of an effective size of N-1. In this case, it was 95 and 94 examples respectively. A receiver operating characteristic (ROC) curve was used to evaluate the cross-validation procedure, that is, the performance of the classifier when faced with new data. An ROC curve graphically expresses the behaviour of the true positive rate versus the false positive rate when the classification threshold (see next paragraph) is varied, as shown in Figure 2. A performance metric derived from the ROC, the area under the curve (AUC) ^26^, is simply the area under a ROC curve. A typical AUC value for a random classifier whose features do not predict the outcome is 0.5, whereas a classifier where the features completely explain the phenomenon would have an AUC of 1.0.

The output of the classifier is the probability *p* that a given example belongs to the RM group. It is possible to set a classification threshold, that is the value of *p* above which an example is considered to belong to the RM group. The true positive rate (sensitivity) and the false positive rate (1-specificity) of the classifier depend therefore on the chosen classification threshold, as setting a low threshold reduces specificity but increases sensitivity and vice-versa.

### Data analysis pipeline

In summary, the algorithm IMMATCH consists of the following steps:

1. Create a data matrix D of size 95×9 (9 features for each of the 95 couples) with the features described in the previous section, and a label vector **I** of size 1×95 with the labels (control or RM) of each couple.
2. Create a submatrix D_i_ of size 94×25 by removing the i-th row (a whole couple) of D, and a sub-vector **I**_i_ (of size 1 ×94) by removing the *i*-th element of **I**. Neither D_i_ nor l_i_ contain any information about the couple *i*.
3. Train an SVM with D_i_ and use it to predict the probability *p* (or score) that couple *i* belongs to the RM group.
4. Repeat the process for all the couples *i*, to form a vector of 95 scores. Compute the AUC with the vector of scores and the label vector **I**.

Steps 1 to 4 are illustrated in Figure 4.

### Statistical significance

A random permutation test was used to estimate statistical significance ^27^. A p-value can be defined as the probability of obtaining an effect at least as extreme as the one observed when the null hypothesis is true. In this case an effect at least as extreme means an AUC value at least as large as the one observed, and the null hypothesis is that the features are not predictors of RM. To estimate this probability, a permutation test shuffles the label vector l to disrupt any possible correlation between the features and the labels (making the null hypothesis hold) and computes the resulting AUC. The previous process was iterated 2000 times, and the *p* value was calculated by computing the fraction of the times that the obtained AUC was equal or greater than the AUC obtained with the non-shuffled vector I. A similar approach was taken to compute the *p* value associated with the accuracy.

### Code availability

The code used for this analysis is available upon reasonable request but it is not publicly available. Requests to access the codes will be reviewed by Immune Compass LTD to verify whether restrictions due to Intellectual Property apply.

### Data availability

The data that support the findings of this study are available upon reasonable request, subject to the agreement of the study participants to share their genotype and clinical data.

## AUTHOR CONTRIBUTIONS

A.M.S, D.I.A.S and I. N participated in the design of the study and the manuscript writing; A.M.S. and D.I.A.S. contributed to the analysis and interpretation of the data.; A.M.S. created the software used in this work.

