## Supplemental Information for "Towards a Gamete Matching Platform: Using Immunogenetics and Artificial Intelligence to Predict Recurrent Miscarriage"

The proposed functions for assessing allele similarity rely on structural, evolutionary and functional properties of the proteins, which are long chains of amino acids.

FUNCTIONS f_1_ and f_2_: CONTINUOUS DISTANCE METRICS FOR ALLELE SHARING

The similarity of any two amino acids can be assessed via a similarity matrix. One example of a type of similarity matrix is the group of BLOSUM matrices ^17^, in which the similarity of two amino acids is assessed as a function of how often one substitutes another in highly conserved regions of protein families. Amino acids with similar properties are therefore more likely to appear at the same site. Another example of similarity matrix is the PMBEC matrix ^18^, which focuses specifically on peptide-MHC binding interactions. The extent to which two residues are considered similar depends on the similarity of their contribution to binding free energy in different environments ^18^.

In general, with any similarity matrix, the distance between two proteins can be computed as a function of the amino acids in which they differ. For this work, the distance between proteins was computed with the MATLAB function *seqpdist*, using the default method (Jukes-Cantor) and either the BLOSUM50 matrix (for function f_1_) or the PMBEC matrix (for function f_2_).

FUNCTION f_3_: NON-BINARY SIMILARITY ASSESSMENT FOR ALLELE SHARING VIA LINEAR ALGEBRA AND PHYSICOCHEMICAL DATA

The unusual diversity of HLA proteins is thought to be driven by selective pressure ^9^, in which case the high cardinality (at each locus, the number of possible proteins is in the order of thousands) would provide adaptive advantages to the population. As mentioned in the introduction, to make the analysis feasible, researchers often represent HLA proteins in ultra-low resolution, for instance, clustering thousands of proteins into two groups. Mathematically, performing such operation is called cardinality reduction. The method presented in this section was motivated by the need of performing efficient cardinality reduction of HLA proteins. Ideally, a functional way to perform cardinality reduction should capture as much of the information of protein diversity while making the mathematical problem tractable. As protein-peptide binding affinity is at the core of HLA proteins' biological functions, we decided to represent each HLA protein as a small set of continuous variables reflecting the binding affinity of said protein to reference peptides. Such representation allows to:

- Have a compact, yet rich representation of the protein as a small set of continuous numbers (The advantages of this will be discussed further in this same section); and
- Use the above set of numbers to assess similarity between two proteins.

The cardinality reduction process that we used was inspired by collaborative filtering for recommender systems. Briefly, a recommender system predicts user preferences for items by using historical data of an ideally large number of users ^20^. A popular approach consists of building a matrix with users as rows and items as columns, or vice versa. The element *(i,j)* represents how satisfied the *i-*th user is with the *j*-th item. Within this formulation, the goal of the recommender system is to predict the matrix’s missing values, that is, the unknown preferences. Researchers then may use a modified version of singular value decomposition (SVD, a linear algebra technique), called incremental SVD, to predict the missing values ^19^. Let us consider *n* users and *m* items: the incremental SVD solves the problem by finding an optimal decomposition of the original matrix into the product of an *n*×*r* and an *r*×*m* matrix, where *r*, the rank of the matrices, is a parameter to be chosen.

Once these two matrices are found, each user and each item thus become a vector of *r* elements, and following the matrix multiplication rules, any missing ranking a*_i,j_* _,_ can be expressed as the following inner product (where **q**_i_ is the vector representing user *i* and **p**_j_ is the vector representing item *j*):

a*_i,j_* = **q**_i_^T^ **p**_j_  …………(1)

While the details of the SVD approach are out of the scope of this work, it is worth stating a few properties that are highly relevant for our problem. First of all, by finding the optimal decomposition, users and items become feature vectors that reflect the structure of the original matrix in a compact manner. Each of the *r* dimensions can be seen as an abstract trait learned by the optimisation process (minimising the error between predictions and known ratings). An item with a high value of a specific dimension means that this trait is evidently representative of the item, and a user with a high value of that dimension means that the user appreciates that trait. In that manner, the product of equation 1 is high when the user *i* displays preference for the traits that the item *j* possesses. The abstract traits learned by the optimisation process not only reflect user preferences in a compact fashion, but they are ordered according to their relevance to data explanation.

An analogy with the HLA proteins problem can be easily drawn, wherein users become HLA proteins, items become reference peptides, and rankings become binding affinities. The *r*-dimensional vector that represents the HLA protein will be referred to as the physicochemical continuous representation of said protein. Binding affinity values were obtained from the Immune Epitope Database ^28^ website as IC50 values. Instead of raw IC50 values, affinity was expressed as -1/log(IC50) to better scale the data. For each HLA protein, the number of IC50 data points (measured binding affinities to reference peptides) ranged from 1 to 29963, with a mean of 2371. For proteins with less than 10 measured IC50 values, the closest protein was used instead, to obtain more robust representations. Protein closeness was assessed using the function f_1_. It is worth mentioning that the motivation behind applying the recommender system analogy is not to predict unknown binding-affinity values but to be able to represent a protein with continuous features based on its physicochemical activity.

The above method can be used for two different purposes. The first is to use the vector of *r* elements directly as a set of features representing the physicochemical activity of the HLA protein. The abstract traits, or the elements of the vector, are already ordered according to their relevance, setting *r* thus would imply setting a cutoff value to the desired number of features. Therefore the value of *r,* that is the dimension of the vector, can be seen as a hyper-parameter of the model. In addition, if a protein can be mapped to a vector, then for any two proteins, the inner product between the two corresponding vectors can assess the similarity of the two proteins, which would be the second purpose. The geometry of the problem is out of the scope of this text, however, it can be insightful to remember that in the case of recommender systems, equation (1) predicts the missing ratings by computing the inner product between a user vector and an item vector. For assessing similarity between proteins, *r,* the dimension of the vectors, can be taken as large as desired, as the inner product will be a scalar regardless the dimension of the vectors. We took *r* to be 500 for this particular implementation. Furthermore, as we are interested in allele sharing, we did not use the vectors as features, we used their values only as intermediate steps to compute the inner products corresponding to the similarity assessment. More specifically, for any two given proteins, f_3_ is defined as the inner product of their physicochemical continuous representations.

The implementation of the incremental SVD used here was a modified version of the Julia package developed by Aaron Windsor (https://github.com/aaw/IncrementalSVD.jl).

As a closing remark for this section, we can further elaborate on the motivation behind choosing a small set of continuous features. Let us assume that we can represent each protein, say HLA-B, as a set of three continuous features. By using three continuous features to represent an HLA-B protein, an individual can be represented by six numbers at that locus (given the two copies of the gene) and a couple can therefore be represented by 12 numbers. As a categorical variable, HLA-B has 3700 possible values. Hence, considering the four copies of a couple, there are 3700^4^ combinations. It might not look like an improvement to change one variable for 12 variables. However, using dummy encoding for instance, 3700^4^ variables would be required to represent one single categorical variable with 3700^4^ categories. On the other hand, a binary assessment allows for the description of two different properties, whereas there are infinite ways to fill a three dimensional space, and therefore, a large number of different behaviours can be represented. The proposed compact description not only dramatically reduces the required amount of data but allows us to exploit a property of continuous numbers, the ability to judge proximity. A representation with continuous features introduces the possibility of assessing how close the predictors of different examples (couples) are and allows an algorithm to infer the outcome based on examples featuring spatially close predictors. Closeness is a key concept here, as it is meaningless for categorical variables. Therefore, such local extrapolation available for continuous variables further reduces the number of examples required.
